# Impact of domain knowledge on blinded predictions of binding energies by alchemical free energy calculations

**DOI:** 10.1101/150474

**Authors:** Antonia S J S Mey, Jordi Juárez Jiménez, Julien Michel

**Affiliations:** EaStCHEM School of Chemistry, University of Edinburgh, David Brewster Road, Edinburgh EH9 3FJ, United Kingdom, E-mail

**Keywords:** D3R, computer-aided drug design, protein-ligand interactions, alchemical free energy calculations

## Abstract

The drug design data resource (D3R) consortium organises blinded challenges to address the latest advances in computational methods for ligand pose prediction, affinity ranking, and free energy calculations. Within the context of the second D3R Grand Challenge several blinded binding free energies predictions were made for two congeneric series of FXR inhibitors with a semi-automated alchemical free energy calculations workflow featuring the FESetup and SOMD tools. Reasonable performance was observed in retrospective analyses of literature datasets. Nevertheless blinded predictions on the full D3R datasets were poor due to difficulties encountered with the ranking of compounds that vary in their net-charge. Performance increased for predictions that were restricted to subsets of compounds carrying the same net-charge. Disclosure of X-ray crystallography derived binding modes maintained or improved the correlation with experiment in a subsequent rounds of predictions. The best performing protocols on D3R set1 and set2 were comparable or superior to predictions made on the basis of analysis of literature SARs only, and comparable or slightly inferior, to the best submissions from other groups.

## 1 Introduction

There is growing interest in the routine use of alchemical free energy (AFE) calculations for predictions of protein-ligand binding energies in structure-based drug discovery programs [1–7]. In particular building on pioneering work over three decades ago [8, 9], some modern alchemical relative free energy calculation protocols achieve in several diverse protein binding sites sufficiently accurate predictions of binding energies (root mean square deviations (RMSD) under 1.5 kcal·mol^−1^; Pearson Correlation coefficient’s (R) of around 0.7 or better) to speed up hit-to-lead and lead optimisation efforts [10]. In favourable cases, AFE calculations can even reproduce subtle non-additivity of structure-activity relationships [11]. However, for a given set of protein-ligand complexes it remains difficult to anticipate the predictive power of AFE calculations. Uncertainties in binding modes [12–14] protonation/tautomeric states [15, 16], binding site water content [17–19], and choice of potential energy functions [20, 21], can profoundly influence the outcome of such calculations. Accordingly, there is much interest in defining as much as possible a domain of applicability for the technology [22].

Blinded prediction competitions, whereby participants submit physical properties computed by a model in the absence of knowledge of the actual experimental data, have been instrumental in driving methodological progress in a wide range of scientific fields [23–26]. Blinded predictions reduce the impact of unconscious biases on the design of protocols, and allow evaluation of molecular modelling methods in a context closer to their intended use in drug discovery. This is advantageous for academic groups that have expertise in computational methodologies, but lack resources to carry out prospective studies. It is also beneficial for the field to evaluate different methodologies applied to the same dataset with identical analysis protocols.

This report focuses on the predictions submitted by our group within the context of the second Drug Design Data Resource (D3R) Grand Challenge, that ran between September 2016-February 2017. The D3R Grand challenge 2 was the second blinded prediction challenge organised by the D3R consortium in this case looking at predicting binding poses, binding affinity ranking, and free energies for a series of 102 ligands of the Frasenoid X Receptor (FXR). This complements previous reports from our group on blinded predictions of protein-ligand poses, rankings, binding free energies [27], distribution coefficients [28], and host-guest binding free energies [29], within the frame work of the first D3R challenge in 2015 and the SAMPL5 challenges [30, 31]. The dataset of 102 inhibitors of FXR, both crystal structures and affinity data, were provided by Roche. The competition featured pose predictions, dataset rankings, and relative binding free energy predictions for two subsets of 15 and 18 compounds referred to as set1 and set2 respectively. Our group only submitted predictions of the relative binding free energies for the set1 and set2 subsets. Submissions were made before (stage1) and after (stage2) information about binding poses of representative set1 or set2 compounds were made available. This enabled an analysis of the impact of the available experimental data on the performance of the protocols. All input data download and submissions upload were conducted via the website of the D3R consortium [32].

## 2 Theory and Methods

### 2.1 Datasets

#### 2.1.1 Blinded datasets

At the start of the challenge (stage1), the organisers released the pseudo apoprotein structure of ligand **10** as provided by Roche, as well as 36 ligands in SDF format to be used for the prediction of crystallographic poses, and an additional set of 66 ligands that should be used in affinity rankings. There were two subsets identified among these 102 ligands, set1 with 15 compounds and set2 with 18 compounds, for which relative binding free energies could be calculated. Ligand subsets set1 and set2 are depicted in Figure SI2. For the second stage of the challenge, 36 X-ray structures were released, meaning that they could be used to prepare input files for alchemical free energy calculations. Once the competition was over a set of IC50 data for the entire dataset was released. The data stems from a scintillation proximity assay using only the FXR binding domain and a radioactive tracer. More information on the experimental binding assay as well as a study on other FXR inhibitors can be found in a series of publications [33–36]. Experimental relative binding free energies were estimated by eq. 1

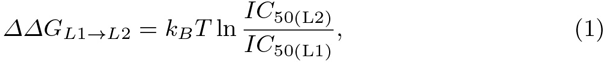

where L1 and L2 represent two ligands for which a relative energy difference is computed and *k_B_* and *T* are the Boltzmann constant and temperature respectively.

#### 2.1.2 Literature datasets

In order to test the computational protocols before submission of blinded predictions, retrospective studies were carried out using available literature data. A set of inhibition and structural data for 3-aryl isoxazole analogs of the non-steroid agonist GW4064 had been previously published [34, 36]. The data consists of two different ligand series, where the first series contains 8 compounds (LitSet1) and the second series 17 (LitSet2). The same experimental IC50 assay as described for the blinded dataset was used. Relative binding free energies were computed from the reported IC50s with equation 1. A summary of the compounds present in LitSet1 and LitSet2 can be found in the Figure SI1.

### 2.2 Methods

The methodology used for the calculations of relative binding free energies of FXR ligands was a single topology molecular dynamics alchemical free energy approach. Several operations are necessary to produce, from an input set of protein atom coordinates and 2D descriptions of ligands, a set of output relative free energies of binding. Currently this is implemented by a semi-automated workflow as depicted in figure 1.

**Fig. 1.**
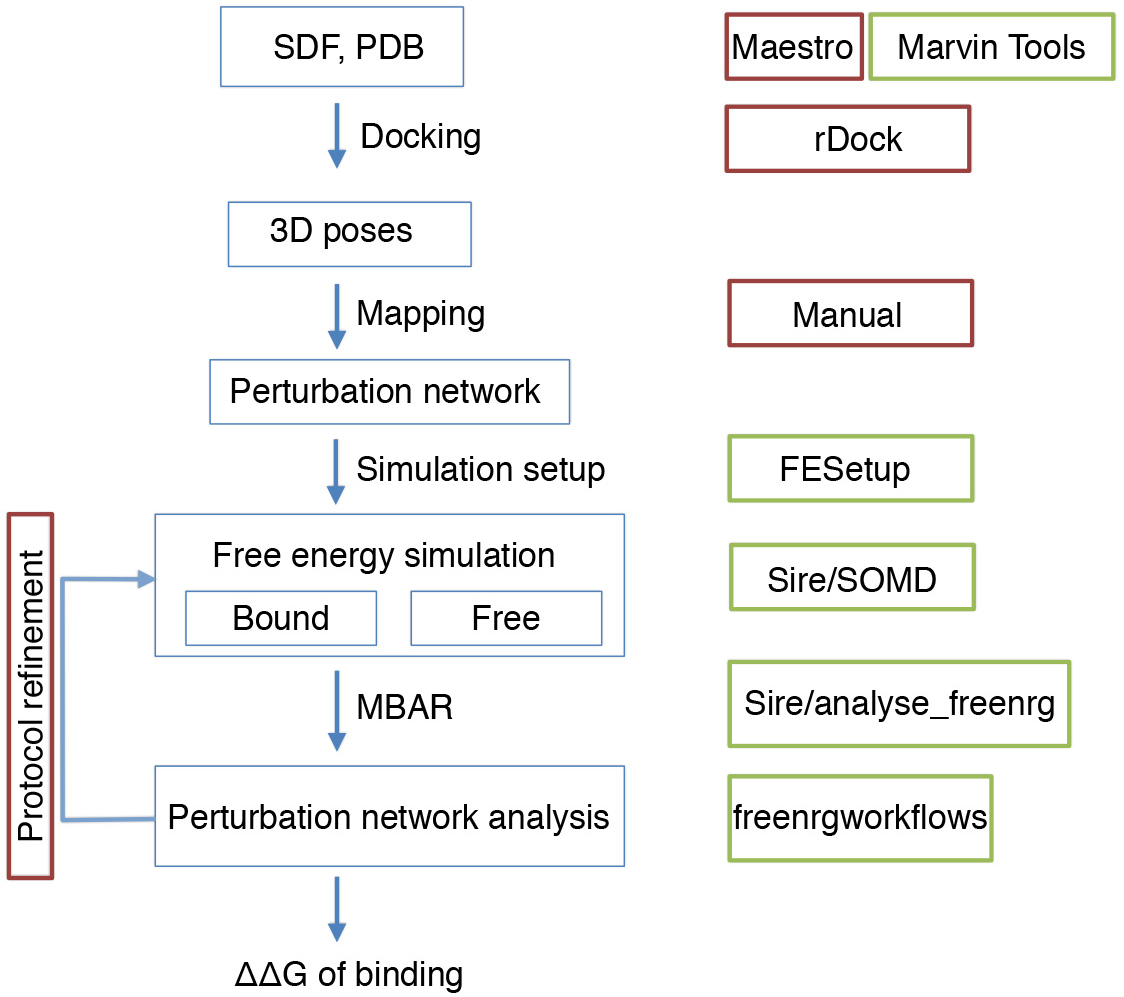
Semi-automated workflow for predicting relative free energies of binding. Workflow operations are depicted by blue boxes. Green boxes denote software available for automated execution of the workflow step. Red boxes denote operations that require human intervention.

#### 2.2.1 Initial protein and ligand structure setup

For the two sets of literature data, the crystal structure with PDB ID 3FXV (FXR in complex with compound **7a**) was used for the ligands taken from Feng et al. [34], and the crystal structure with PDB ID 3OKI (FXR in complex with compound **1a**) was used for data taken from Richter et al. [36].

Due to the plasticity of the binding site of FXR and the differences in shape between compounds in set1 and set2, two different protein structures were needed to build complexes between FXR and compounds of set1 and set2. Each structure required a different preparation protocol. For set1 the FXR structure provided by the organizers was chosen as an initial template. For the docking calculations, that mainly consider residues delineating the binding site, the standard protein preparation workflow in Maestro 11 (beta) and conversion to the appropriate format with the utility fconv was sufficient. To use the resulting structure in alchemical free energy simulations, however, it was necessary to model the missing region comprised between residues A459 and K464. Visual analysis of crystallographic structures available in the PDB revealed that fragments of the region comprised between M450 and N472 are missing in several structures (i.e: 3FXV), or are arranged in at least two slightly different conformations. The first conformation displays a slightly kinked alpha helix spanning from residue N432 to residue N461 with a loop connecting residues D462 to T466 (as in structure 3OKH). In the second conformation the kinked alpha helix is shorter (N432 to S457) and the loop is longer (W458 to T466) and adopts a different orientation (as in structure 3OKI). After superimposing the structure provided by the organizers with representative structures of each conformation, 3OKH was deemed as a suitable template to build the missing fragment of the structure. Subsequently appropriate capping groups were added to residue M247 of the main chain and to residues D743 and D755 of the co-activator fragment. For set2, the 3OKI structure was used as an initial template and the preparation process was significantly simpler. The standard protein structure preparation workflow of Maestro 11 (beta) with addition of capping groups was sufficient to generate structures suitable for both docking and FEP calculations.

Ligand 3D structures compatible with the assay conditions were generated from 2D sdf files provided by the organizers using MarvinTools scripts available in Marvin Sketch 15.3.30 software package. The pKa predictor available in the same package was used to evaluate the major protomer/tautomer for these compounds bearing ionizable substituents. No crystallographic water molecules were retained for the docking calculations.

#### 2.2.2 Generation of ligand binding modes

Binding modes for the literature data were manually build in Maestro 11 (beta) by means of an overlay with the binding mode of compounds **7a** and **1a** as observed in the X-ray crystal structures 3FXV and 3OKI respectively.

For set 1 of the blind datasets, a putative binding mode for the series was obtained by docking the compounds bearing the smallest (hydrogen, **91**) the bulkiest (morpholino amide, **102**) substituent, as well as compound **101** to probe the effect of a ioniozed carboxylic acid on the binding mode. Consistent binding modes were obtained for the three molecules in the crystallographic structure provided by the organizers. To minimize the differences between binding modes within the set1 series, the binding modes for all compounds were manually created from the binding mode of the largest compound **102**. A similar protocol was followed for compounds in set2, using compounds **12**, **74**, **76**, **79** and **83** to explore the influence of different substituents in the sulfonamide. A consistent binding mode was found for these compounds in the protein conformation displayed in PDB ID 3OKI, and putative binding modes for the entire series were manually created from the binding mode of compound **83**.

All docking calculations were performed with rDock, generating the cavity using the two-sphere method available in the program, centering a 15 Å cavity within residues M294, I356, S336 and Y373 using 1.5 and 4.0 Å for the radius of the small and large spheres respectively. Manual building of the compounds was performed with Maestro 11 (beta) and the minimizer available in the suite was used to avoid steric clashes. After poses were obtained, water molecules resolved in the X-ray structure provided by the organizers were superimposed with the coordinates of the poses. Clashing water molecules were displaced to nearby positions. For the second stage of the challenge, the additional knowledge gained from the crystal structures was leveraged to prepare new input files for the alchemical free energy calculations.

#### 2.2.3 Alchemical calculations input preparation

Once a set of satisfactory 3D poses for both set1 and set2 was obtained, a relative free energy perturbation network was manually designed for both set1 and set2 ligands. The network was assigned in such a way that resulting perturbations between structure would be minimal and as many as possible simple cycles would be contained in the network in order to allow for cross validation using cycle closure as a measure. Set1 included one ambiguous binding mode for compound **47**. For set2 only three of the 18 compounds had a clearly preferred binding mode. Typically there was uncertainty in the position of ortho or meta substituents of a benzyl ring. Whenever there was ambiguity, the different binding modes were included in the perturbation map. The perturbation networks can be found in figures 3-7 of the SI. With the perturbation networks defined, the software FESetup [37] release 1.3dev, was used to parametrise set1 and set2 ligands, setup ligands in a water box as well as protein environment and create the needed input for the alchemical free energy simulations.

##### Ligands

Ligands were parametrised using the generalised amber force field 2 (GAFF2) [38], followed by solvation in a rectangular box of 12 Å length using TIP3P water [39, 40]. An energy minimization using a steepest decent algorithm with 500 steps was carried out on the water box, followed by an NVT simulation with the ligand restrained, during which the system was heated to 300 K over 1000 steps. Next an NPT equilibration at 1 atm was run for 5000 steps, followed by the release
of the restraint on the ligand over 500 steps. FESetup used the software pmemd for this equilibration. For each perturbation a SOMD compatible perturbation file was then created from the perturbation map produced by FESetup.

##### Protein ligand complex

For the protein and ligand complex the protein and previously parametrised ligands were combined and solvated in a rectangular box of 10 Å. The protein forcefield was the amber 14 SB forcefield [38]. An equivalent solvation and equilibration protocol was used as described for the solvated ligand only.

#### 2.2.4 Alchemical free energy simulations

The alchemical free energy protocol used here is based on the SOMD software as available in the Sire 2016.1.0 release [41]. This version of SOMD is linked with OpenMM 7.0.1 [42] that provides a CUDA compatible integrator enabling simulations to be run on a cluster of GPUs.

Details about the theoretical background are available elsewhere [6, 43, 7, 4, 44– 46, 10]. The main idea behind alchemical free energy calculations is to avoid direct computation of the free energy change associated with the reversible binding of a ligand to a protein. Instead one computes the free energy change for artificially morphing a ligand (*L*_1_) into another ligand (*L*_2_). By introducing a parameter *λ*, which defined the change from *L*_1_ to *L*_2_. Practically, either a replica exchange algorithm is used to simulate at different *λ* windows, or a set of discrete *λ* simulations is carried out. Repeating this process for *L*_1_ and *L*_2_ in aqueous solution or bound to the protein of interest enables construction of a thermodynamic cycle that yields the relative binding free energy of the two ligands.

Each alchemical free energy calculation for a pair of ligands *L*_1_ and *L*_2_ consisted minimally of one forward (*L*_1_ to *L*_2_) and one backward (*L*_2_ to *L*_1_) computation. Ligand pairs that showed poor agreement between forward and backwards simulation were repeated up to three times. Mean free energies and standard error were estimated from the resulting distributions of computed relative binding free energies. Further details are provided in the SI [47]. All simulations shared the following common set of parameters. Each simulation box was treated with periodic boundary conditions and simulations were run for 4 ns each using a 2 fs integration timestep with a Leap-Frog-Verlet integrator. Bonds involving hydrogens were constrained, except if the hydrogen atom was morphed to a heavy atom in the perturbation. The temperature was maintained at 298 K using an Andersen thermostat and a collision frequency of 10 ps^−1^ with velocities initially drawn from a Maxwell-Boltzmann distribution of that temperature. Pressure was kept at 1 atm using the Monte Carlo Barostat implemented in OpenMM with an update frequency of 25 MD steps. For non-bonded interactions an atom-based shifted Barker-Watts reaction field scheme was used with a cutoff of 10 Å and the reaction field dielectric constant *ε* = 82.0. The number of *λ* windows for each simulation varied for different perturbations and a summary, as well as complete simulation parameters can be found in the SI. All input files are available on a github repository [48].

#### 2.2.5 Free energy analysis and convergence

Free energy changes were estimated both with thermodynamic integration and the multi state Bennett’s acceptance ratio (MBAR) estimator, as implemented in pymbar (v 3.0.0 beta 2) [49], which was integrated into the Sire app *analyse_freenrg*. The TI analysis served mainly two purposes; first, to ensure that the MBAR and TI free energy estimates for a particular perturbation are consistent to within approximately 0.5 kcal·mol^−1^ and second, to test the convergence of the gradient 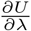 time series. Stationarity of the timeseries was assessed by means of the augmented Dicky-Fuller test. Non-stationary gradient timeseries in the pool of over 10,000 timeseries trajectories generated in this study were identified. This served as a basic test for convergence, and all non-stationary trajectories were repeated. Convergence was also assessed by checking whether binding free energies from forward and backward simulations were consistent, as well as with cycle closures in the perturbation network. Simulations with poor cycle closures or poorly agreeing forward and backward transformations were repeated multiple times. The actual process of the free energy analysis for estimating cycle closure and overall affinities based on MBAR is described in the following. The first 5% of the trajectories were discarded to allow for equilibration before MBAR analysis. Perturbation for morphing *L*_1_ to *L*_2_ and *L*_2_ to *L*_1_ were both simulated and resulting binding free energies were averaged for the forward and (reversed) backward perturbations. When available, averages were calculated across multiple independent repeats. The individually estimated free energy differences were then read into a networkx (v 1.11) digraph [50]. The error estimated between repeated runs of backwards/forwards simulations served as the estimated error for each averaged network edge. Binding free energies relative of a ligand *L_i_* to a reference compound *L*_0_ were then estimated by enumerating all possible simple paths connecting *L_i_* to *L*_0_ in the network. The relative binding free energy and its uncertainty along a given path was obtained by summing relative binding free energies along each edge of the path and propagating errors. A simple path in a network is defined as the path between two vertices *v_p_* and *v_q_*, with no vertex repeating along the path. Therefore a path between ligand *L_q_* and *L_p_* can be written as *P_p,q_* = (*v*_1=*p*_*, v*_2_, …, *v*_*n*=*q*_). This path is only valid if every pair of vertices has an entry in the weighted adjacency matrix (*w_ij_*), which in this cases holds the free energy difference of each perturbation. Therefore, the relative free energy along a single simple path with *n* vertices, is be given by: Δ*g_p,q_* = *w*_*p,*2_,+…+*, w*_*n−*1*,q*_. The associated error of the path can be obtained from the error matrix *ϵ_ij_*, which similarly to the weighted adjacency matrix will hold the error associated with each edge in the network. The error for a given simple path is therefore given by: 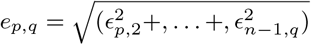. Based on the error associated with each simple path a weight of the path can be defined as 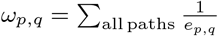. Therefore the relative free energy between ligand *L_p_* and *L_q_* can be defined as the weighted average of all paths, using *ω_p,q_* as the path weight.

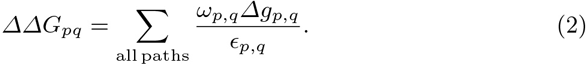

The corresponding error *E_p,q_* to the estimated free energy is give by:

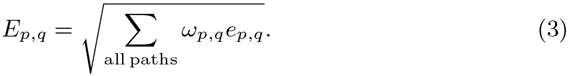

Thus paths that have smaller statistical errors contribute more than paths that show larger statistical errors.

If multiple binding modes for one compound were used in the network, they were combined into a free energy for a single compound using exponential averaging in the following way:

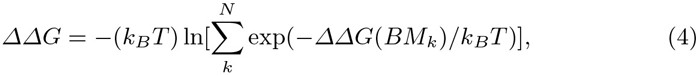

where *k_B_* is the Boltzmann constant, *N* the total number of binding modes, and *BM_k_* denotes the k-th binding mode.

#### 2.2.6 Charge scaling correction

Initial analysis of literature datasets (see results) suggested that polarisation effects may play a significant role in FXR ligand binding energetics. While no polarisable force-field was readily available to test this hypothesis, there has been some success in capturing polarisation effects in protein-ligand binding by QM/MM reweighting of trajectories computed with a classical potential energy function [51]. Given the time constraints posed by the competition, no such methodologies were used here. Rather, an *ad hoc* protocol based on empirical scaling of ligand partial charges was implemented.

Thus the corrected free energies were given by:

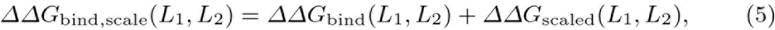

where *ΔΔG*_scaled_ is given by:

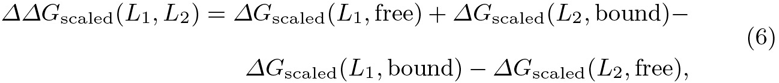

where the *ΔG_scaled_* values are the free energy changes for scaling the partial charges of a ligand *L*_1_ or *L*_2_ in water, or bound to FXR. Such quantities were evaluated via MBAR analysis of trajectories for 5 evenly spaced *λ* windows sampled for 1 ns each. Scaling factors ranged from 1 (no scaling) to 0.5 (50% decrease in magnitude of partial charges).

### 2.3 Errors analysis

For the comparison of computed binding free energies and experimental binding free energies two measures are mainly used in the analysis, Pearson R and mean unsigned error (MUE). To obtain an error estimate for both these measures a bootstrapping approach is used in which the mean and standard error of each of the computed free energy estimates serve as the mean and standard deviation of a Gaussian distribution. For each estimate a new value from this Gaussian distribution is drawn until a new artificial distribution of computed free energies is sampled. This resampled distribution is then correlated to the experimental data. Repeating the process 10,000 times gives rise to a distribution of MUE and R, for which a mean and 1*σ* confidence interval can be computed. This was the default protocol used to estimate metric errors. The organisers, however, chose a different way of estimating errors in the data sets to facilitate comparison between different submissions. This approach uses bootstrapping of the dataset, for which data points (both experimental and computed) are resampled with replacement until an artificial dataset of the same size is created. This is repeated 1,000 times, leading to a distribution for Pearson R with 1*σ* confidence intervals. All error bars in figure 6 have been generated in this fashion.

## 3 Results

### 3.1 Literature datasets

The robustness of the computational protocol was first tested with the two literature datasets LitSet1 and LitSet2. Supplementary figures 3 and 4 depict the perturbation network used for LitSet1 and LitSet2 respectively [47]. A summary of the results, comparing the calculated and measured binding free energies is given in table 1. While the correlation between LitSet1 computational and experimental data with R=0.84±0.05 was deemed satisfactory, the mean unsigned error (MUE) at 3.0±0.2 kcal·mol^−1^ was judged unexpectedly large. For the second dataset LitSet2 the overall correlation R=0.56±0.03 is lower, however the MUE is significantly lower at 1.7±0.1 kcal·mol^−1^.

**Table 1.** Summary of test dataset based on GW4064 compounds. *Charged compound **1R** has been omitted from the analysis.

| Dataset | scaling factor | R | MUE / kcalomol<sup>-1</sup> |
| --- | --- | --- | --- |
| LitSet1 | 1.0 | 0.84 ± 0.05 | 3.0 ± 0.2 |
|  | 0.9 | 0.83 ± 0.04 | 2.45 ± 0.18 |
|  | 0.8 | 0.81 ± 0.04 | 2.24 ± 0.23 |
|  | 0.7 | 0.78 ± 0.08 | 1.8 ± 0.15 |
|  | 0.6 | 0.56 ± 0.09 | 2.2 ± 0.2 |
|  | 0.5 | 0.51 ± 0.1 | 1.4 ± 0.2 |
| LitSet2* | 1.0 | 0.56 ± 0.05 | 1.77 ± 0.08 |
|  | 0.9 | 0.54 ± 0.05 | 1.54 ± 0.09 |
|  | 0.8 | 0.51 ± 0.05 | 1.46 ± 0.08 |
|  | 0.7 | 0.44 ± 0.06 | 1.47 ± 0.09 |
|  | 0.6 | 0.37 ± 0.06 | 1.61 ± 1.7 |
|  | 0.5 | 0.23 ± 0.07 | 1.75 ± 0.08 |

Analysis of the pairwise alchemical free energy calculations in the two datasets suggested that calculated binding free energy changes for perturbations that involve substitution of a non polar group by a polar group were overly exaggerated with respect to experimental data. Also, LitSet2 contained one negatively charged compound (**1R**, carboxylic acid) which was predicted to be ca. 30 kcal·mol^−1^ less potent than its -H counterpart, whereas experimental data suggests weaker binding of the acid by ca. +2.6 kcal·mol^−1^.

The binding site of FXR is rather apolar (see figure 2A), and it was hypothesized that changes in ligand polarisation upon transfer from bulk to the FXR binding site may play a significant role. This prompted the development of an *ad hoc* protocol in an attempt to capture polarisation effects as described in the methods section via introduction of a set of charge scaling factors. The resulting correlation coefficient and MUE for scaling factors between 1 to 0.5 are also displayed in table 1. Figure 8A in the SI displays the correlation between the computed and experimental results, and figure 8B of the SI summarises the effect of changing the scaling corrections from 0.9 to 0.5 of the original charge. It was found that a scaling factor of 0.7 was the best tradeoff to minimize MUE whilst maintaing a reasonable Pearson correlation coefficient. The effects are more pronounced for LitSet1. The one exception is the charged compound **1R** in LitSet2, for which reasonable agreement with experimental data required a scaling factor of 0.5.

**Fig. 2.**
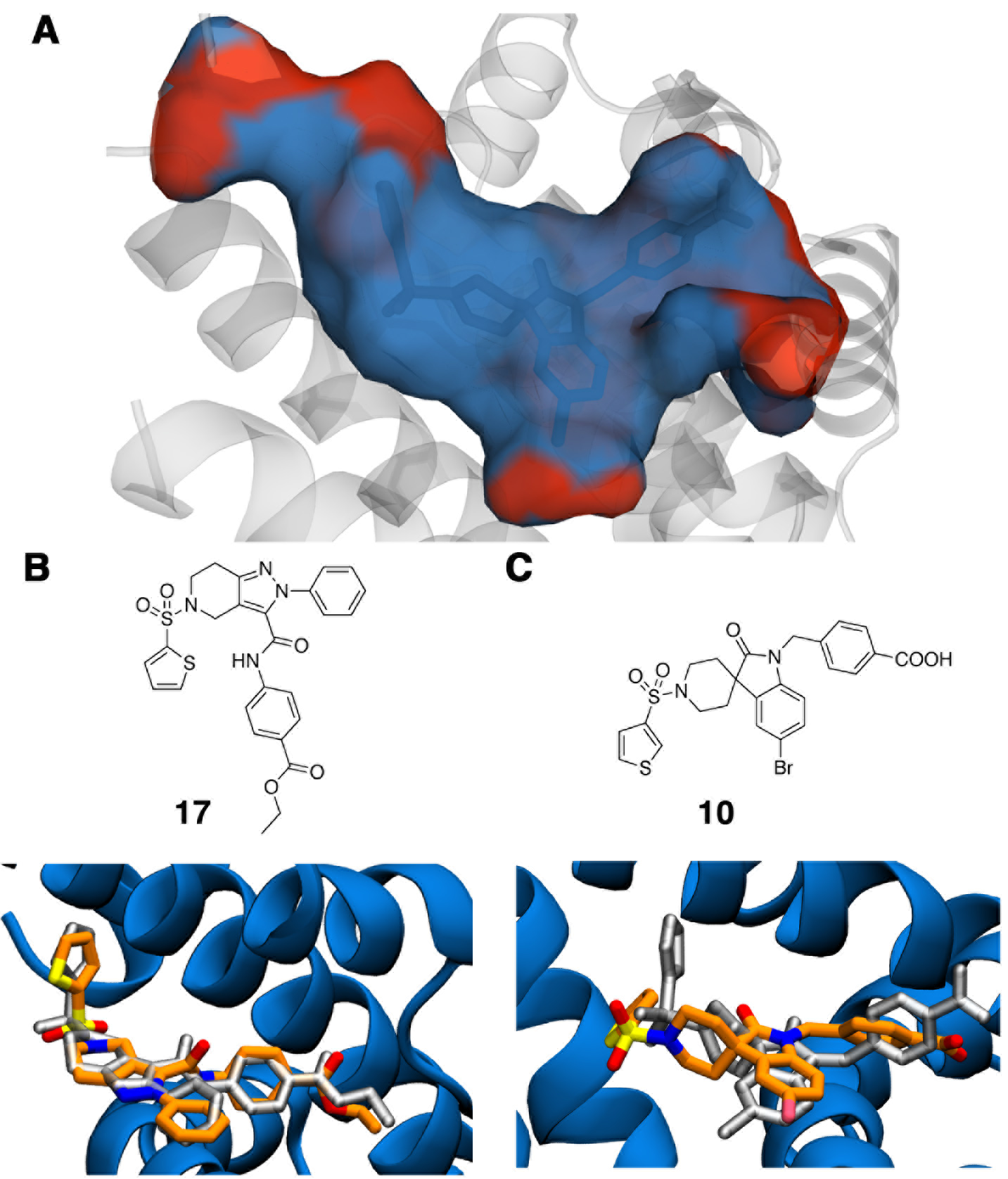
A: Depiction of the FXR binding site, with hydrophilic residues shown in red and hydrophobic residues shown in blue. B: Compound **17** from set1, and predicted (orange sticks) versus observed (grey sticks) binding modes. C: Compound **10** from set2, and predicted (orange sticks) versus observed (grey sticks) binding modes.

Given time-constraints no further efforts were devoted to the literature datasets, and subsequent blinded submissions were made for protocols without charge scaling correction, or with a charge scaling correction of 0.7 for free energy perturbations that maintain net-charge, and 0.5 if the net-charge varies in the perturbation.

### 3.2 Blinded dataset

For the first stage of the competition binding modes for the FXR ligands in set1 and set2 had to be predicted by analysis of available crystal structures, or docking calculations as described in the methods section. Figure 2B top panel shows the structure of set1 ligand **17**, and the bottom panel depicts predicted and later disclosed binding modes. The RMSD is 0.9 Å only, and the binding mode prediction can be considered successful. For set2 the X-ray crystal structure of **10** was later disclosed. The predicted binding mode deviates more, whereas at 2.5 Å the RMSD is not exceptionally high, the thiophene ring has been positioned differently to the X-ray pose. This was of concern as many of the set2 compounds feature variations in aryl sulfonamide groups.

Table 2 shows results for the protocols submitted at stage 1 of the competition. The *expert opinion full* protocol was a submission where binding energies were predicted by one of the authors (JM) by analysis of literature structure activity relationships and visualisation of predicted binding modes for the LitD1 and LitD2 datasets. The *expert opinion same charge* protocol was not submitted but is presented to facilitate comparison with other protocols. The *full* protocol was a submission on the full dataset analysed as described in methods. The *full guided* protocol was a submission where only a small number of pathways in the perturbations network were hand-picked by one of us (JM) to evaluate binding free energies for the dataset. This was only done for set1. Finally the *same charge* protocol was a submission of alchemical free energy predictions restricted to the largest subset of compounds with the same net-charge.

**Table 2.**
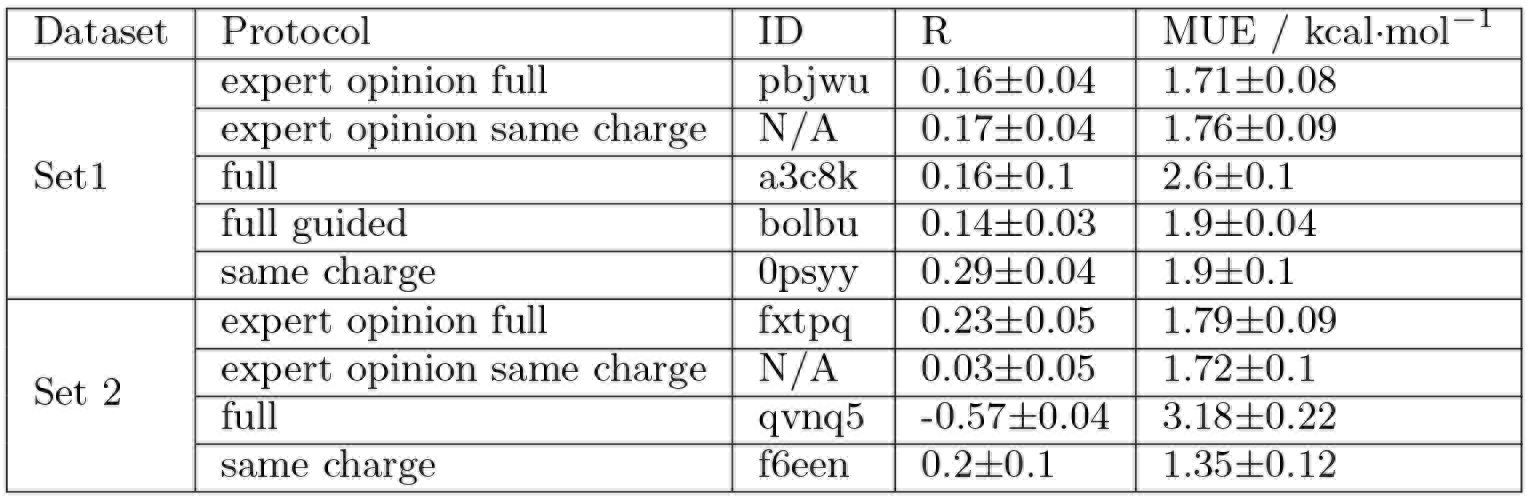
Performance of the protocols submitted at stage 1 of the D3R competition.

For the second stage of the competition, calculations were repeated from a new set of poses for set2 compounds. set1 poses were the same as in stage 1. Additionally individual perturbations were categorised as ‘easy’, ‘medium’ or ‘difficult’ on the basis of the precision of the calculated relative binding free energies obtained at stage 1, and this led to lambda schedule protocols with less, the same amount of, or more, windows as in stage 1 (see SI for details). The time left in the competition was used to carry out multiple repeats of the perturbations that showed higher statistical errors. Additionally, the optimisation of charge scaling factors on the literature datasets had been completed by then, and scaling factor corrections were also applied to set1 and set2 datasets. Table 3 shows the results for protocols submitted at stage 2 of the competition. Only *full* and *same charge* protocols, and their scaled variants, were submitted.

**Table 3.**
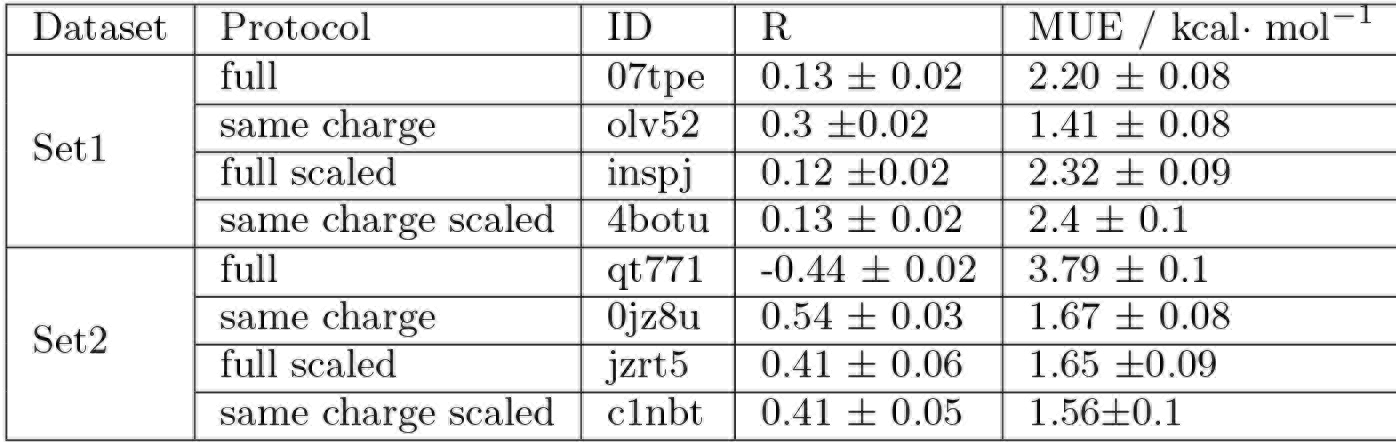
Performance of the protocols submitted at stage 2 of the D3R competition.

At stage 1 the *expert opinion* protocol shows R values for both set1 and set2 of ca. 0.2, and MUE values ca. 1.7 kcal· mol^−1^. The performance is similar or worse for the *expert opinion same charge* protocol. Alchemical free energy based protocols on the full dataset fare poorly with similar or lower R values, and higher MUE values. Submissions that only considered compounds with the same net charge show better performance (R ca. 0.2-0.3, MUE ca. 1.4-1.9 kcal· mol^−1^). Overall none of the protocols show satisfactory correlation with experiment.

At stage 2 of the competition, the *full* and *same charge* submissions show lower statistical errors because the additional repeats calculations on the noisier perturbations have improved convergence. For set1 the MUE decreases, but the R metric is no different from stage 1 submissions. The scaled submissions for the full dataset and the same charge dataset achieve similar R values, but the MUE has worsened. For set2 lower statistical errors are also observed with respect to stage 1. The *full* submission produces similarly low R values and high MUE values. However the *full scaled* submission significantly increases R from ca. -0.4 to +0.4, while decreasing MUE from ca. 3.8 to 1.6 kcal· mol^−1^. The *same charge* submission shows a significant increase in R with respect to stage 1 (from ca. 0.2 to ca. 0.5), but the MUE increases from 1.3 to 1.7 kcal· mol^−1^. Finally, the *same charge scaled* protocol achieves a poorer R value (ca. 0.4) and similar MUE value.

Overall the most significant improvement at stage 2 is observed for the set2 same charge dataset. This could be because the predicted binding modes at stage 1 were not in agreement with the subsequently disclosed X-ray structures. The scaling protocol appears to yield large improvements on set2, but this actually comes at the expense of decreased predictive power for the subset of compounds that carry the same net-charge (see below).

Figure 3 depicts detailed results for the full dataset of set1 compounds at different stages of the competition. Figure 3A shows at stage 1 the relative binding free energy of charged compound **101** is significantly overestimated with respect to all other neutral compounds. Compound **45** is also a significant outlier. Figure 3B shows that at stage 2 there is a trend towards better agreement with experiment, apart from **101** and **45** that remain significantly off. Figure 3C shows that the *full scaled* submission considerably improves free energy estimates for compound **101**, but also drastically decreases the accuracy of estimates for neutral compounds **102, 48, 95, 96**. It is not clear why predictions for **45** consistently perform so poorly.

**Fig. 3.**
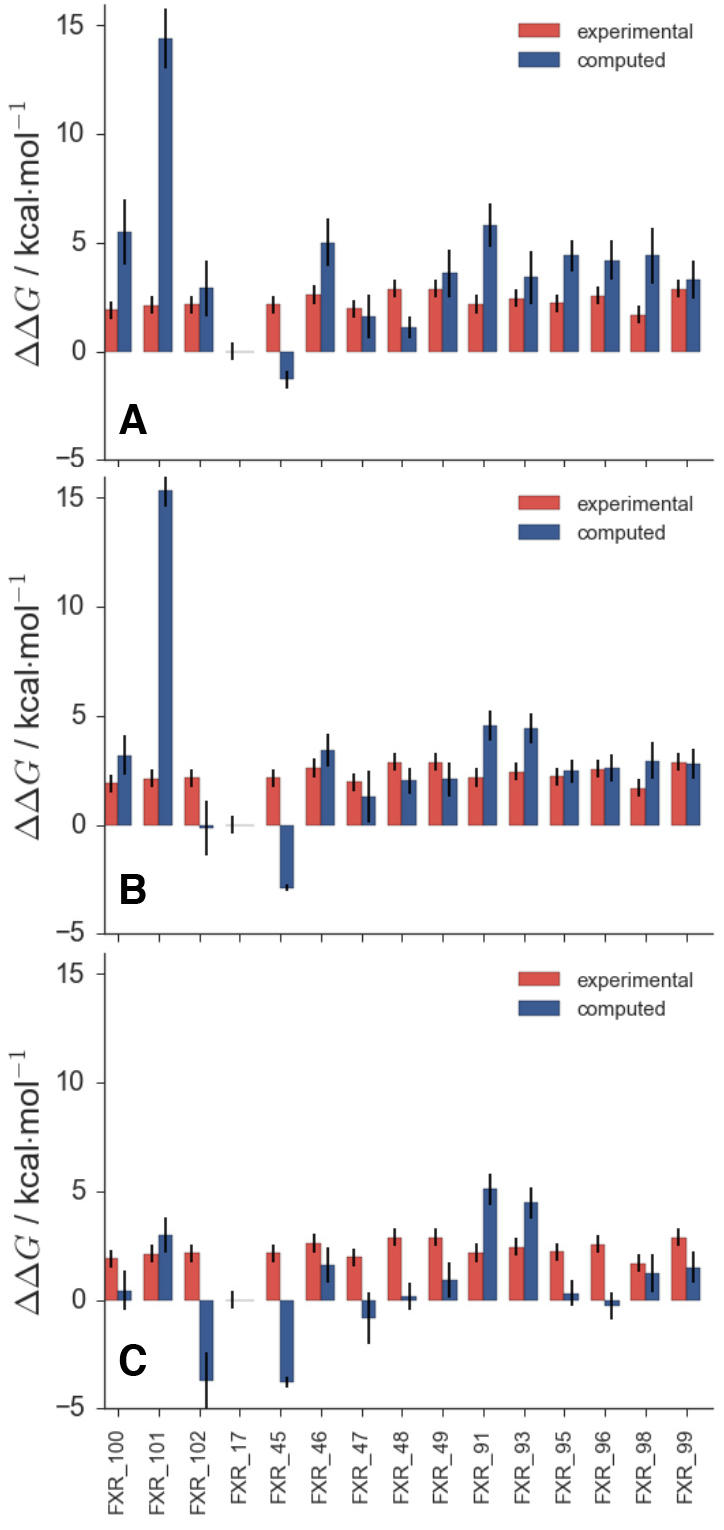
A: Stage1 *full* submission for set1(ID a3c8k) showing clear overestimation of the relative binding free energy of charged compound **101**. B: Stage 2 *full* submission (ID 07tpe). C: Stage 2 *full scaled* submission (ID inspj).

Some highlights for the binding free energy estimations of set2 are shown in figure 4. The stage 1 *full* submission is depicted in figure 4A. The poor estimation of the relative binding free energy of neutral compounds (**38, 41, 73, 75**) with respect to other compounds in the dataset that carry one negative charge is very apparent. Figure 4B, shows the *full* submission at stage 2 of the competition. Estimates for the charged compounds improve, however the binding free energies of the neutral compounds are still poorly estimated. This is more apparent by inspection of figure 5, which shows the *same charge* submission where only charged compounds were considered. This is the best performing protocol overall in terms of correlation coefficient of R = 0.54 ± 0.03 and MUE = 1.67 ± 0.08 kcal· mol^−1^.

**Fig. 4.**
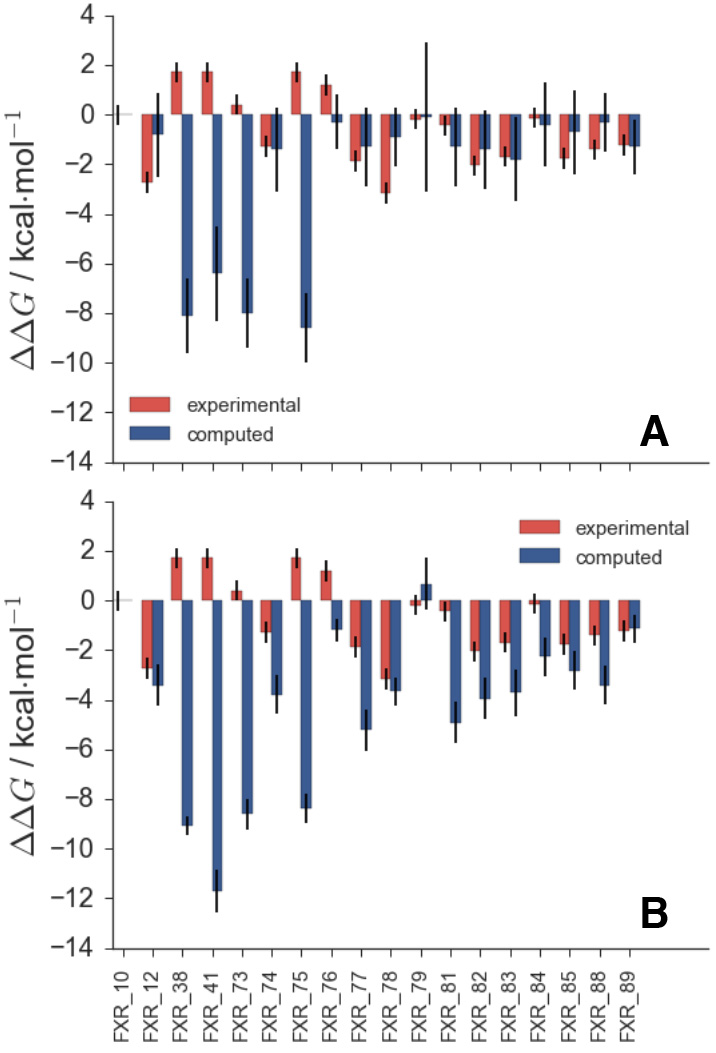
A: Stage 1 *full* submission for set2 compounds (ID qvnq5). B: Stage 2 *full* submission for set2 compounds (ID qt771).

**Fig. 5.**
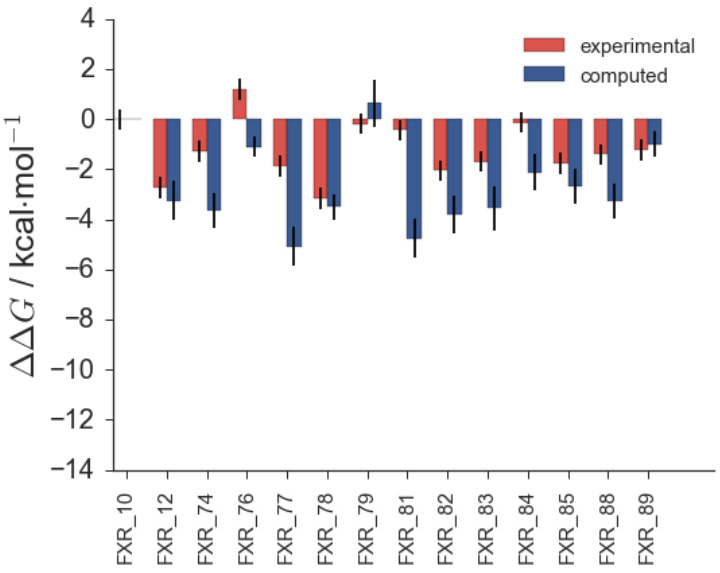
Stage 2 *same charge* submission (ID 0jz8u).

### 3.3 Comparison to other submissions

The organisers released data for all binding free energy prediction submissions shortly after the end of stage2, and a summary of the correlation coefficients can be seen in figure 6. Figure 6A and B are stage 1 submissions for set1 and set2 respectively. Results for stage 2 set1 and set2 are shown in figure 5 C and D. The authors submissions are shown in green and can also be identified by their submission ID listed in table 2 and table 3. It should be noted that the shown correlation coefficients are slightly different to the ones reported in the tables 2 and 3. This is down to the use of different error analysis methods between the authors and the organisers, as discussed in the methods section 2.2. What becomes apparent however, from figure 6 is that for set1 both at stage 1 and stage 2 there is no protocol that obviously outperforms another protocol and no statistically significant ranking is possible. For set2 and particularly stage 2 there are four protocols that perform better than the rest, which are a mix of alchemical free energy and other protocols. Submission **81n55** is an alchemical method based on FEP+, submission **xk67c** uses a non-equilibrium pulling approach using Gromacs as the simulation framework, submission **67a3e** uses the software MIX to perform energy minimisation of protein-ligand complexes, and submission **x2j7p** also used FEP+.

**Fig. 6.**
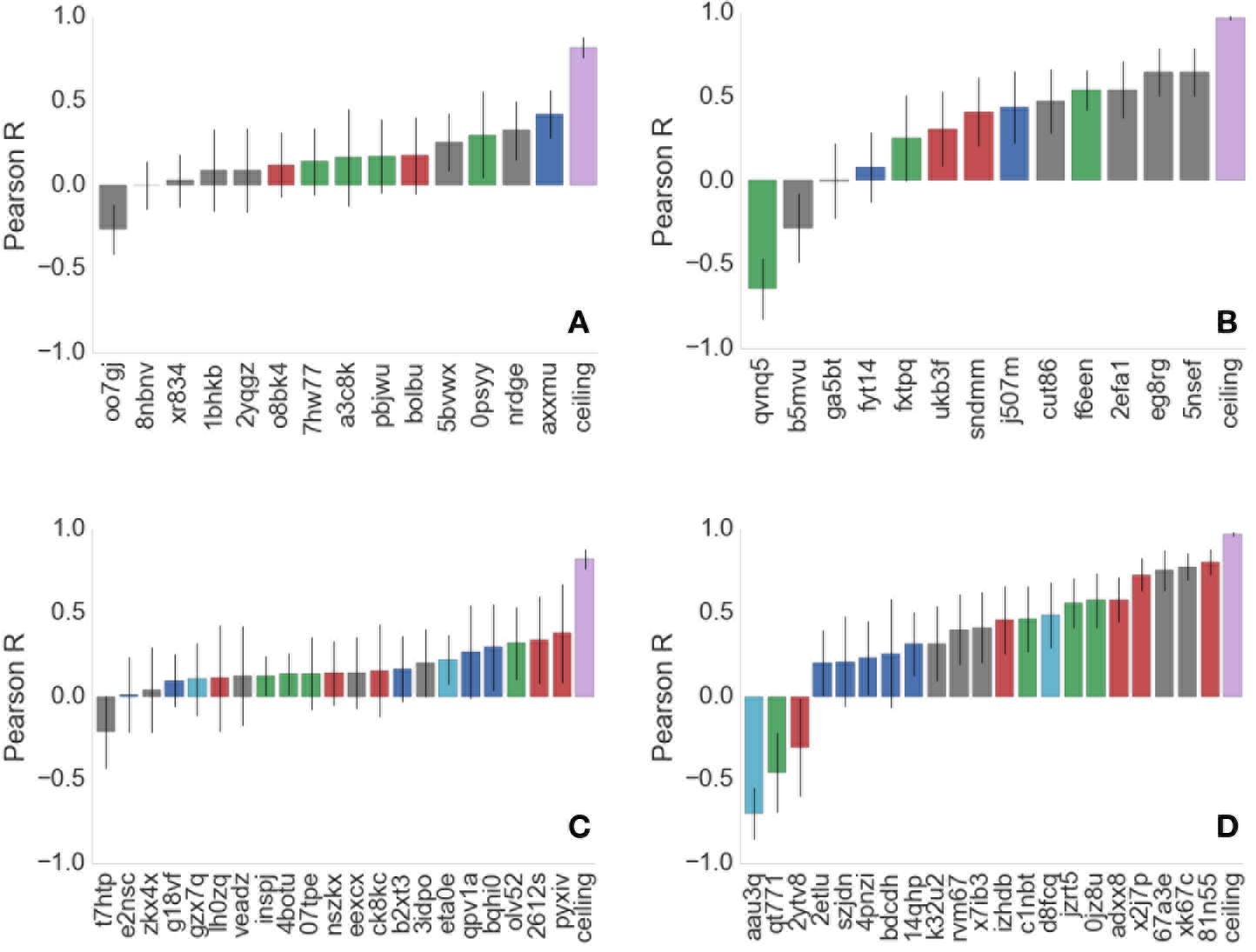
Summary of all submitted protocols. A: Stage 1 for set1. B: Stage 1 for set2. C: Stage 2 for set1. D: Stage 2 for set2. Green colours denote the authors submissions and protocol IDs can be identified in table 2 and table 3. Red colours denote other alchemical methods, blue colours denote MMPBSA based methods, the light blue colour denote quantum mechanical based methods and grey denotes any other methods. The ceiling entry is discussed in the text and shown in purple. All method descriptions are made available by the organisers and can be found on the www.drugdesigndata.org website.

Figure 6 suggests, that there is no clear trend indicating a method that consistently outperforms others. Furthermore, the overall correlation between predicted free energies and experimental values is poor and unreliable for set1.

## 4 Conclusions

The blinded predictions on FXR ligands highlighted difficulties in reliably estimating relative binding energies between compounds that differ in their net-charge with the current workflow. This was anticipated in light of past experience and until methodological advances lift this limitation, alternative ad-hoc protocols may prove more reliable. For instance, other groups submitted in this competition alchemical binding energy predictions for ligands modelled as protonated acids in order to maintain the same net-charge across the full D3R datasets. While this is an unlikely chemical state for the unbound or bound ligands given the assay conditions, this setup did lead to superior predictions for the full D3R datasets. The relatively reasonable correlations obtained retrospectively on LitSet1 and LitSet2 ligand series were encouraging, but the high mean-unsigned error on the LitSet1 dataset prompted the development of an approximate charge scaling protocol to account for potential neglect of polarisation effects. This had no beneficial effect on the accuracy of the blinded predictions and this protocol is not recommended for further use. Overall this indicates difficulties in reliably anticipating the robustness and transferability of the protocol across different ligand series, let alone different binding sites. In spite of the difficulties encountered it is useful to note that expert opinion based on analysis of literature SARs proved no more predictive on set1 and worse on set2 (excluding charged compounds). While this observation lacks statistical relevance – presumably there would also be variability in different expert opinions – it does highlight the difficulty of the problem. By contrast, expert opinion often fares well for poses prediction when compared against automated software workflows [27, 52]. It was also encouraging that the correlation for set2 same-charge subset increased once experimental data about the binding mode of a representative set2 ligand could be taken into account. By contrast no significant variation was observed for set1 upon repeating the calculations, presumably because the binding modes had been well predicted at stage 1 of the competition.

The D3R Grand Challenge 2016 free energy datasets were markedly larger than those used in the 2015 competition. This enabled a more reliable comparison of the performance of different methodologies. Nevertheless, it is apparent that both set1 and set2 are still too small to reliably rank a large number of submissions made by different groups. A general trend for alchemical free energy protocols can be observed, establishing them typically in the top 33% of submission, in particular in set2, for both correlation coefficient and root mean square error (RMSE), as shown in the SI. It is noteworthy that the features of the distribution of experimental binding energies for set1 (shorter span and uneven density) contribute to making predictions intrinsically more difficult than for set2. In addition, the precision of the experimental data was not determined. Assuming a ca. 0.4 kcal·mol^−1^ uncertainty in experimental measurements [53] together with bootstrapping suggests ceiling values for R of ca. 0.82±0.06 and 0.97±0.01 for set1 and set2 respectively. Thus the best performing methods are far from achieving high accuracy R values on set1, but show respectable correlation on set2. While it may be difficult to source significantly larger datasets amenable to alchemical free energy calculations, it may be useful to assess their intrinsic difficulty for the design of future competitions.

## Acknowledgements

J.M. is supported by a Royal Society University Research Fellowship. The research leading to these results has received funding from the European Research Council under the European Unions Seventh Framework Programme (FP7/2007-2013)/ERC grant agreement No. 336289. J. J-J is supported by a Marie Sklodowska-Curie fellowship grant agreement No 655677.

